# Experimental infections with a honeybee virus induce fitness costs in ant colonies

**DOI:** 10.1101/2025.10.22.683975

**Authors:** Lai Ka Lo, Ashmita Baruah, Robert J. Paxton, Yuko Ulrich

## Abstract

Acute Bee Paralysis Virus (ABPV) is a key driver of honey bee colony losses that has been increasingly reported in non-bee hosts. Ants have long been hypothesised to act as viral reservoirs, but most evidence comes from field surveys, and experimental tests are still scarce. Here, we combined survival and transmission assays, viral load quantification, and viral replication assays following experimental inoculations to test whether the clonal raider ant *Ooceraea biroi* can harbour and transmit ABPV within its colonies. ABPV-injected immatures and adults showed delayed development and elevated mortality, respectively. Viral replication assays suggested these fitness costs were caused by host responses (e.g., immunopathology) rather than viral replication. Viral particles persisted for days in inoculated ants and high viral loads were detected in untreated colony members after 1–3 days of cohabitation with ABPV-injected pupae or adults, indicating rapid viral spread. These results show that ants can acquire ABPV, incur fitness costs, and pass the virus within their colonies, suggesting that they may act as incidental viral reservoirs. By maintaining and disseminating honey bee viruses, even without supporting replication, ants may contribute to their environmental persistence and spillover across species.

## 1. Introduction

Honey bees are vital pollinators for ecosystems and agriculture, yet highly vulnerable to human activities and environmental changes. Over the past two decades, substantial winter colony losses of western honeybees, *Apis mellifera*, have been reported worldwide (Neumann & Carreck, 2010). Several iflaviruses and dicistroviruses have emerged as major drivers of colony losses in both managed and wild populations (Carreck et al., 2010; Genersch & Aubert, 2010; McMenamin & Genersch, 2015; Neumann & Carreck, 2010; Yañez et al., 2020). While these viruses can persist covertly at low titres without causing apparent symptoms, the introduction of the ectoparasitic mite *Varroa destructor* (henceforth: varroa) has facilitated viral spread and turned asymptomatic infections into lethal diseases (Wiegers, 2021) by injecting viruses into the bee haemolymph (Ryabov et al., 2014).

Acute Bee Paralysis Virus (ABPV), a +ssRNA virus in the family *Dicistroviridae* (Benjeddou et al., 2001; de Miranda et al., 2010), has been detected repeatedly in collapsing and *Varroa*-infested colonies and is considered a significant driver of honey bee mortality (Ball, 1985; Ball & Allen, 1988; Berthoud et al., 2010; Cox-Foster et al., 2007; de Miranda et al., 2010; Žvokelj et al., 2020). It occurs across all host life stages and castes, with pupae being especially permissive to viral replication (Azzami et al., 2012; Bailey & Milne, 1969; de Miranda et al., 2010; Žvokelj et al., 2020). Experimental infections have shown that the injection of fewer than 100 viral particles into pupae or adults can induce tremors, paralysis, and death within days (Bailey et al., 1963). ABPV has been detected outside honeybees (Doublet et al., 2025; Gusachenko et al., 2020; McMahon et al., 2015; Pascall et al., 2021) and is considered a multi-host virus, with cross-species transmission facilitated by shared floral resources and nesting sites (Yañez et al., 2020).

In recent years, ants have been proposed as viral reservoirs for ABPV based on repeated field detections (Celle et al., 2008; Gruber et al., 2017; Levitt et al., 2013; Lin et al., 2020; Payne et al., 2020; Rivas Fontan et al., 2025; Sébastien et al., 2015). With an estimated 20 quadrillion individuals and a biomass exceeding that of all wild birds and mammals, ants are globally dominant insects (Parker & Kronauer, 2021; Schultheiss et al., 2022) whose foraging and nesting ranges overlap with those of many taxa, making them potential pathogen vectors (Dobelmann et al., 2023; Tiritelli et al., 2025). ABPV has been reported in ants and honey bees from ant-infested hives, suggesting ants as viral vectors within apiaries (Rivas Fontan et al., 2025). Yet, most studies remain observational, and whether the virus can persist, transmit, and replicate in ant, is largely unknown. Experimental work is limited to one study, which showed that oral exposure to ABPV impaired locomotion and colony growth in *Lasius niger* (Schläppi et al., 2020).

Here, we investigate ABPV infections in the clonal raider ant *Ooceraea biroi*, a powerful model for studying disease dynamics under controlled laboratory conditions. Workers in *O. biroi* colonies reproduce clonally and synchronously (Ravary et al., 2006), allowing precise experimental control over genotype and age at the individual and colony levels. In addition, the predatory lifestyle of *O. biroi*, whose diet includes ants and other insects (Kronauer, 2009), makes them prone to pathogen spillover. To investigate the consequences of ABPV exposure in this system, we measured the effects of controlled inoculations on host survival and assessed the ability of the virus to replicate within hosts and transmit within colonies.

## 2. Materials & Methods

### (a) Ant rearing

All ants were sourced from stock colonies of genetic lineage B (Kronauer et al., 2012), and maintained at 28 °C in airtight plastic containers (8 x 12 x 5 cm) with a humidified plaster of Paris floor (Knauf) before experiments. The nests were regularly cleaned, re-humidified, and fed with sterilized, frozen ant brood (*Tetramorium bicarinatum* and *Oecophylla* sp.) and houseflies. For each experiment, all ants were clonally related.

### (b) Infection assay in pupae and adults

#### (i) Preparation of viral inoculum

The ABPV inoculum was prepared as described by Tehel et al. (2019). Briefly, 1 µL of homogenate from a heavily ABPV-infected *A. mellifera* worker was injected into white-eyed pupae obtained from virus-free colonies. The injected pupae were incubated for three to five days to allow viral replication and systemic infection, after which they were homogenised in 0.5 M cold potassium phosphate buffer (PPB, pH 8.0). The resulting homogenates were screened using RT-qPCR to confirm the absence of other common bee viruses, such as deformed wing virus (DWV), black queen cell virus (BQCV), chronic bee paralysis virus (CBPV), Israeli acute paralysis virus (IAPV), sacbrood virus (SBV), and slow bee paralysis virus (SBPV). Only homogenates confirmed to contain ABPV exclusively were used in the experiments. Control inoculum was prepared from uninjected white-eyed pupae processed in the same way.

#### (ii) Inoculation via microinjection

10-day old ant pupae were placed on glass slides and injected with viral inoculum (ABPV at 10^5^ genome equivalents, GEs) laterally at the posterior abdomen using a FemtoJet microinjector (Pi = 200, Pc = 30; Eppendorf AG, Hamburg, Germany) and borosilicate glass capillaries (8.89 cm length, 0.530 mm ± 25 μm inner diameter, 1.14 mm outer diameter; World Precision Instruments). Inoculum was injected until the pupal body visibly stretched. Control pupae were injected with the control inoculum following the same procedure. Following inoculation, eight pupae of either treatment were placed together with six paint-marked, uninjected cohabiting adults in plastic cups (diameter 4.5 cm) with a humidified plaster of Paris floor. For each treatment (control and ABPV-injected) and each sampling time point (0, 3, and 5 days post-inoculation, d.p.i.), four replicate colonies were established. At time point 0, pupae were flash-frozen immediately after injection. At later time points (3 and 5 d.p.i.). Both pupae and cohabiting adults were frozen to assess viral transmission. Pupae survival and development were assessed in the colonies sampled 5 d.p.i. (four control and four infected colonies). Pupae that disappeared, turned dark, or were partly cannibalized were classified as dead.

Adult ants (mixed age) were immobilised by placing them in slits cut in pencil erasers (Läufer Plast 0120) and injected with 10 nL of viral inoculum (ABPV 5 × 10^4^ GEs) or control inoculum at a rate of 5 nL/s under the largest abdominal tergite using a NanoFil 36 GA bevelled needle connected to a NanoFil syringe and an electronic microinjection pump (UMP3 pump with Micro2T SMARTouch controller, World Precision Instruments, USA). Ten injected adults from either treatment were housed with five paint-marked, uninjected cohabiting adults in plastic cups (diameter 4.5 cm) with a humidified plaster of Paris floor. For each treatment (control and ABPV-injected) and each sampling time point (0, 1, 3, and 5 d.p.i.), four replicate colonies were established. At time point 0, focal ants were flash-frozen immediately after the injection. At later time points (1, 3, and 5 d.p.i.), both focal ants and cohabiting adults were flash-frozen. The colonies sampled 5 d.p.i. were monitored for survival over the full period before sampling.

### (c) Viral load quantification

For each colony, five injected adults or three injected pupae were flash-frozen in liquid nitrogen and stored at -80 °C. In the ABPV-injected groups, five cohabiting adults were also sampled. Total RNA was extracted from pooled, whole-body samples using a combined Trizol/phenol-chloroform-column protocol (Libbrecht et al., 2016) and eluted in 20 µL of nuclease-free water. RNA concentration and purity were assessed using a BioPhotometer D30 (Eppendorf, Germany).

Viral load quantification was performed according to a protocol modified from Tehel et al. (2019). For cDNA synthesis, 200 ng of total RNA was reverse-transcribed using oligo(dT)18 primers (Thermo Scientific) and reverse transcriptase (M-MLV and Revertase, Promega, Mannheim, Germany) according to the manufacturer’s instructions in a Veriti Thermal Cycler (Applied Biosystems). The resulting cDNA was diluted 1:3 and stored at -20 °C. qPCR was performed using the KAPA SYBR FAST Universal kit (Kapa Biosystems, Sigma-Aldrich) on a LightCycler 480 (Roche), with two technical replicates per sample, and the following thermal cycle: three min at 95 °C; 40 cycles of 10 s at 95 °C, 30 s at 57 °C, 20 s at 72 °C; final elongation for 1 min at 95 °C; melt curve analysis was performed from 55 °C to 95 °C with 0.5 °C per second increments to check for amplification specificity. Forward primer (5′-CATGGCTCAAGACACTTCATCG-3′) and reverse primer (5′-CCAGCAATGACCTCAATGTG G-3′) were used to amplify a 105 bp fragment specific to ABPV capsid protein gene, were designed with Primer-BLAST (Ye et al., 2012). Each plate included negative controls (control inoculum and no-template control) and a positive control (viral inoculum). Samples with Ct values >35 or with high variation between technical duplicates (Ct standard deviation >0.5) were re-assayed, except for negative controls and control inoculum, with actin (host reference gene) amplified using primers Ob-actinF (5’-GCCGATCGTATGCAGAAGGA-3’) and Ob-actinR (5’-ACCCTTGGCGATGTGTCTTT-3’) designed with Primer-BLAST. To quantify the number of viral GEs in each sample, external DNA standards were generated from the viral inoculum using primers ABPV-F6531 (5′-TCATACCTGCCGATCAAG-3′) (Locke et al., 2012) and ABPV-R6774 (5’-CCAGCAATGACCTCAATGTGG-3’). The PCR product size (244 bp) was verified by electrophoresis on a 2% agarose gel, and the corresponding band was excised and purified using the NucleoSpin Gel and PCR Purification Kit (Macherey-Nagel, Düren, Germany) according to the manufacturer’s instructions. The purified PCR product was quantified using a spectrophotometre and used to generate ten-fold serial dilutions. Viral loads were expressed as log_10_ GEs per individual ant by dividing total GEs by the number of individuals in each sample.

### (d) Negative-sense strand detection

In a separate experiment, three-week-old adult ants were injected with 10 nL of viral inoculum (ABPV 10^4^ GEs), control inoculum, or left uninjected; five focal adults from each treatment were housed with five paint-marked, uninjected cohabiting adults, as described above. A lower dose was chosen to increase survival and extend the observation period. For each treatment and time point (0, 6, 12, 24, and 48 hours post-injection, h.p.i.), four replicate colonies were established. At each time point, focal and cohabiting ants from each replicate were pooled separately, flash-frozen in liquid nitrogen, and stored at -80 °C. RNA was extracted as described above.

Focal and cohabiting ants sampled 24 h.p.i were used to assess viral replication via negative-strand detection. This method relies on RT-qPCR amplification of negative-strand RNA intermediates, which serve as markers of active replication of positive-strand RNA viruses within host cells, rather than the presence of inert genomic RNA (de Miranda et al., 2010; Yue & Genersch, 2005). Two-step RT-PCR was performed using a tagged forward primer designed to specifically anneal to the complementary DNA (cDNA) product of the negative-sense RNA strand of ABPV. Superscript III reverse transcriptase (Invitrogen, USA) was used for cDNA synthesis. In preliminary tests, all ant and bee samples produced non-specific PCR products using previously published primers (Schläppi et al., 2020). To avoid false positives, we designed new primers for the assay. A total of 200 ng RNA was diluted to 10 µL with water and combined with 1 µL of the tagged forward primer tag-ABPV(-)F5185 (5’-AGCCTGCGCACCGTGG-TTGGTTTGGTGCA GAAGGTG-3’) and 1 µL 10 mM dNTPs. The mixture was incubated at 65 °C for 5 min and then placed on ice for at least 1 min. Subsequently, 4 µL of 5X First-Strand Buffer, 1 µL of 0.1 M DTT, and 1 µL of Superscript III Reverse Transcriptase were added, bringing the total reaction volume to 20 µL. The reaction was incubated at 55 °C for 50 min, followed by incubation at 70 °C for 15 min to inactivate the enzyme. cDNA was purified using the NucleoSpin Gel and PCR Clean-up Kit (Macherey-Nagel, Düren, Germany) and eluted in 20 µL of elution buffer. The purified cDNA (diluted 1:3 with nuclease-free water) served as the template for PCR amplification with Taq polymerase (New England Biolabs), using a tag oligonucleotide (5’-AGCCTGCGCACCGTGG-3’) as the forward primer and the corresponding reverse primer, ABPV(-)R5380 (5’-AGAAAAGTCCATAGGCCCGT-3’). PCR reactions without the tag primer were performed for all samples to verify the complete removal of residual tagged forward primers using the following cycling conditions: initial denaturation at 95 °C for 3 min; 35 cycles of denaturation at 95 °C for 20 s, annealing at 56 °C for 20 s, and extension at 68 °C for 30 s; and final extension at 68 °C for 2 min. The resulting PCR products were analysed by electrophoresis on a 2% agarose gel. A 196 bp PCR product was expected in samples containing negative-sense RNA, indicating active viral replication. Bee pupae injected with control inoculum or ABPV inoculum served as negative and positive controls, respectively.

### (e) Replication Statement

| Scale of inference | Scale at which the factor of interest is applied | Number of replicates at the appropriate scale |
| --- | --- | --- |
| Colony | Colony | 4 control-injected colonies, 4 ABPV-injected colonies for all 4 time points |

### (f) Statistical analyses

All data analyses were performed using R version 4.4.1 (R Core Team, 2024). Models were selected based on Akaike’s Information Criterion (AIC) (Akaike, 1976) and compared using the *anova* function to test if reduction of model terms significantly improved model fit. The final model assumptions and residual diagnostics were assessed using the *DHARMa* package (Hartig, 2020). Post-hoc pairwise comparisons were performed with the *emmeans* package (Lenth et al., 2022) with Tukey’s adjustment for multiple comparisons.

#### Survival

Due to complete separation in the pupal development data (no deaths in the control group and no eclosion in the ABPV-inoculated group), Wilcoxon-Mann-Whitney tests were used to compare the proportion of pupae that hatched into adults or died between treatment groups. For adult survival, the data violated the proportional hazard assumption, and endpoint mortality was therefore analysed using a generalised linear model (GLM) with a binomial error structure and logit link, with Treatment as a fixed factor and Colony as a random factor.

#### Viral load

The viral load in focal and cohabiting ants was quantified by converting the Ct values to GEs using a standard curve, followed by a log_10_ transformation. These transformed titres were analysed using a linear model (*lm* function, base R) with one focal and one cohabiting sample per colony. To assess how viral load changed over time in focal individuals, models included Treatment (control-injected vs. ABPV-injected) and Time point (0, 3, and 5 d.p.i. for pupae; 0, 1, 3, and 5 d.p.i. for adults), and their interaction (Treatment × Time point) as fixed effects. To examine the viral load differences between focal and cohabiting individuals, a separate model was fitted for overlapping time points between the two groups including Individual type (focal vs. cohabiting) as a fixed factor, along with Treatment and Time point. For pupae infection assay, comparisons between focal and cohabiting individuals were performed only within the virus-inoculated treatment group as cohabiting adults were only sampled in colonies of virus-inoculated pupae; thus, only Individual type and Time point were included as fixed effects.

## 3. Results

ABPV-injected ant pupae had drastically reduced eclosion rates compared to control-injected pupae 5 d.p.i. (mean ± SD: 0 ± 0% in ABPV, 75 ± 10.2% in controls; Wilcoxon–Mann–Whitney test: z = 2.477, p = 0.029; Figure 1A). Similarly, ABPV-injected pupae had higher mortality compared to controls (mean ± SD: 31.3 ± 16.1% mortality in ABPV, 0 ± 0% mortality in controls; Wilcoxon–Mann–Whitney test; z = -2.461, p-value = 0.0286).

**Figure 1.**
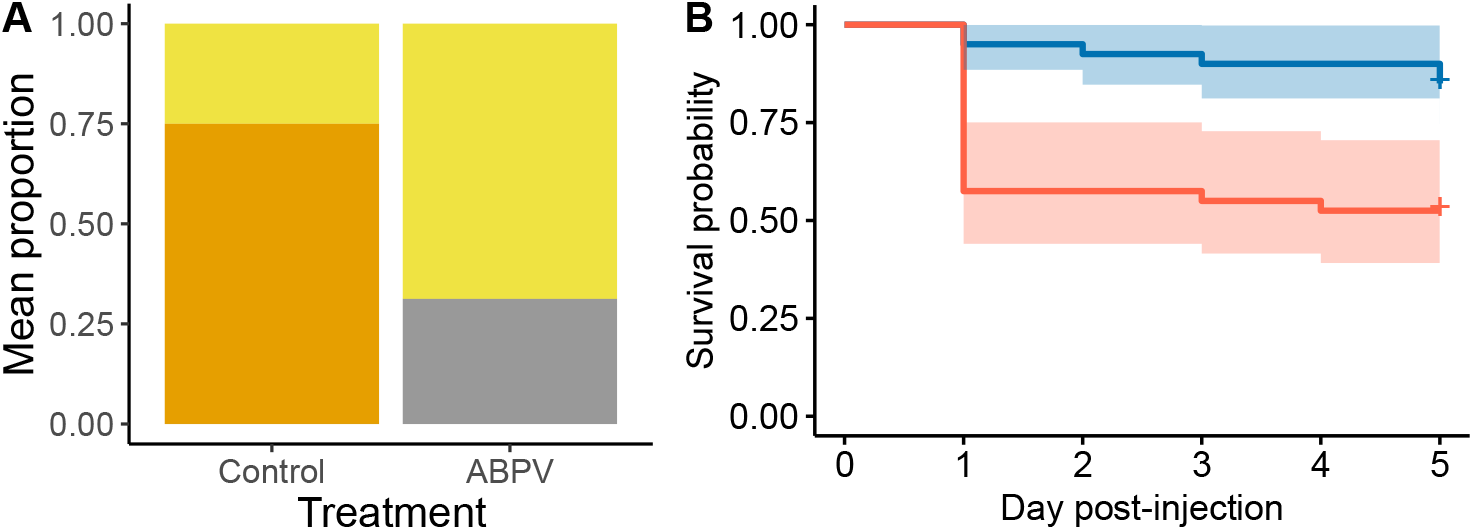
ABPV affects *O. biroi* development and survival. **(A)** Development and survival of ABPV-injected (10^5^ GEs, n=4 colonies) and control-injected pupae (n=4 colonies), shown as the mean proportion of pupae (yellow) that eclosed into adults (orange) or died (gray) five days post-injection. **(B)** Survival of ABPV-injected (5 × 10^4^ GEs, red, n=40 individuals) and control-injected (blue) adult ants (n=40 individuals). Kaplan–Meier curves with 95% confidence intervals.

ABPV-injected adult ants had higher mortality than control-injected adults (mean ± SD: 47.5 ± 31% in ABPV; 15 ± 12.9% in controls; χ^2^(1) = 4.628, p = 0.031), a 5.1-fold increase in the odds of death, mostly occurring 1 d.p.i (Figure 1B). Although virus-injected focal pupae and adults showed increased mortality, no deaths were observed among the cohabiting adults.

Viral load differed between ABPV-injected and control-injected ant pupae (ANOVA: F_1,18_ = 51.802, p < 0.001) and across time points (F_2,18_ = 5.207, p = 0.016) (Figure 2A). As expected, ABPV-injected pupae had higher viral loads than control-injected pupae at all time points (Tukey’s post-hoc; Control vs. ABPV at 0 d.p.i.: t = -7.197, p < 0.0001; 3 d.p.i.: t = -5.726, p <.0001; 5 d.p.i: t = -3.500, p = 0.003). In ABPV-injected pupae, viral load remained consistently high up to 5 d.p.i. (Tukey’s post-hoc; 0 vs. 5 d.p.i.: t = 1.224, p = 0.447).

**Figure 2.**
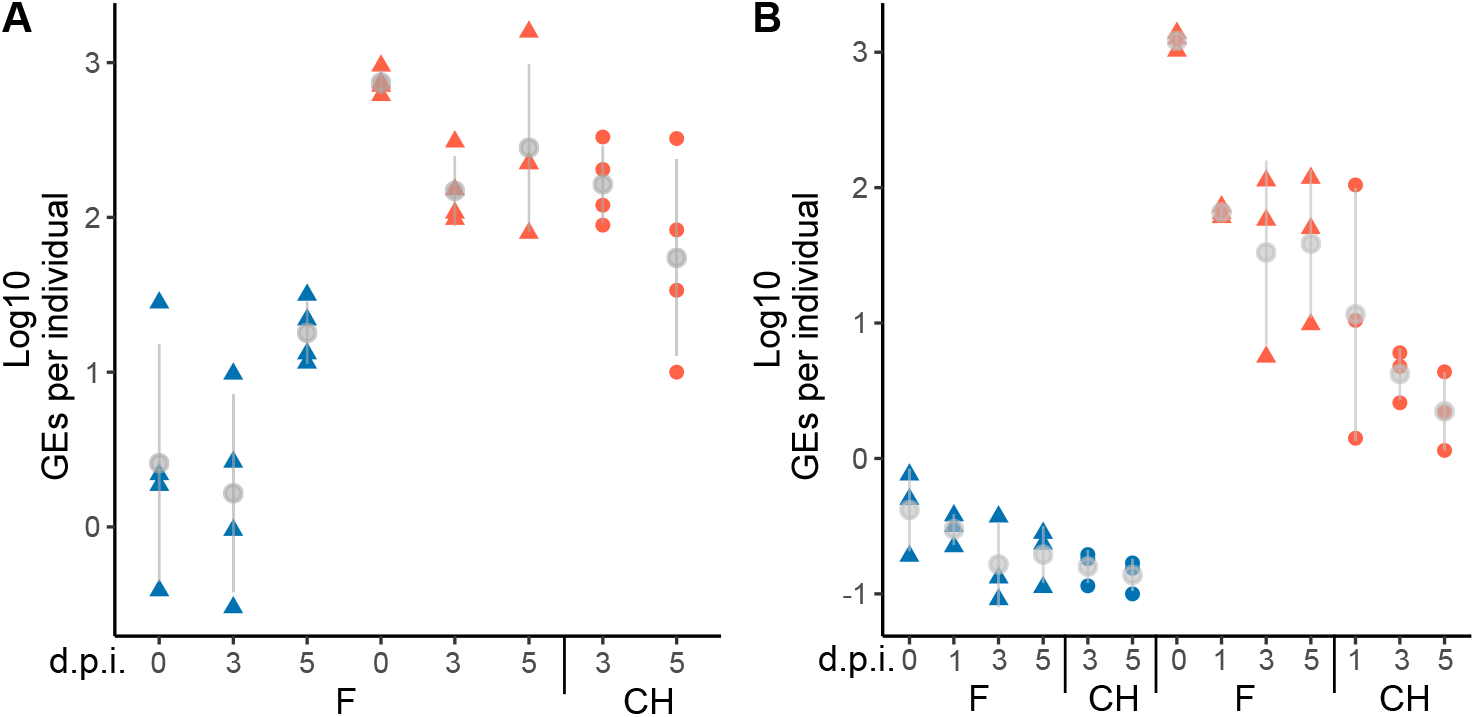
ABPV spreads within ant colonies. **(A)** Viral load in control-injected pupae (blue triangles), ABPV-injected pupae (red triangles), and cohabiting adults (red circles). Data points represent pools of three pupae or five cohabiting adults. Grey: mean ± SD. **(B)** Viral load in control-injected adults (blue triangles), ABPV-injected adults (red triangles), and cohabiting adults (blue and red circles). Data points represent pools of five adults (n = 3-4 per treatment and time point). d.p.i.: days post-injection. Grey: mean ± SD.

In adult ants, viral loads differed between control- and ABPV-injected groups (ANOVA: treatment: F_1,16_ = 140.252, p < 0.001), and this difference varied across time points (ANOVA; treatment × time point interaction: F_3,16_ = 3.87, p = 0.030, Figure 2B). This effect was driven by a sharp decline in viral load in ABPV-injected adults within the first day post-injection (Tukey’s post-hoc; 0 vs. 1 d.p.i.: t = 4.306, p = 0.003), decreasing from approximately 10^3.08^ to 10^1.42^ GEs per individual. Thereafter, viral titres remained at a similar level up to day 5 (Tukey’s post-hoc; 1 vs. 5 d.p.i.: t = 0.795, p = 0.856).

We consistently detected ABPV in uninjected adults that had cohabited with ABPV-injected pupae (Figure 2A) and adults (Figure 2B) for one to five days. Strikingly, adults cohabiting with ABPV-injected pupae carried viral loads as high as those of the injected pupae themselves (ANOVA; focal pupae vs. cohabiting adults, F_1, 13_= 1.922, p = 0.189), suggesting efficient transmission from pupae to adults. In contrast, adults cohabiting with ABPV-injected adults had viral loads lower than those of the injected adults (Tukey’s post-hoc; cohabiting vs. focal: t = -5.381, p <0.0001), yet they still had significantly higher viral loads than controls (Tukey’s post-hoc; control-cohabiting vs. ABPV-cohabiting: t = -6.615, p < 0.0001). This indicates rapid and efficient horizontal transmission of the virus, both from the brood to adults and between adults.

In line with the lack of increase in viral load over time, we found no evidence of viral replication in adult ants. Viral negative-strand replication intermediates were consistently detected in positive controls (ABPV-injected honey bee pupae), but not in ABPV-injected, control-injected, and cohabiting adult ants 24 h post-injection (Figure 3).

**Figure 3.**
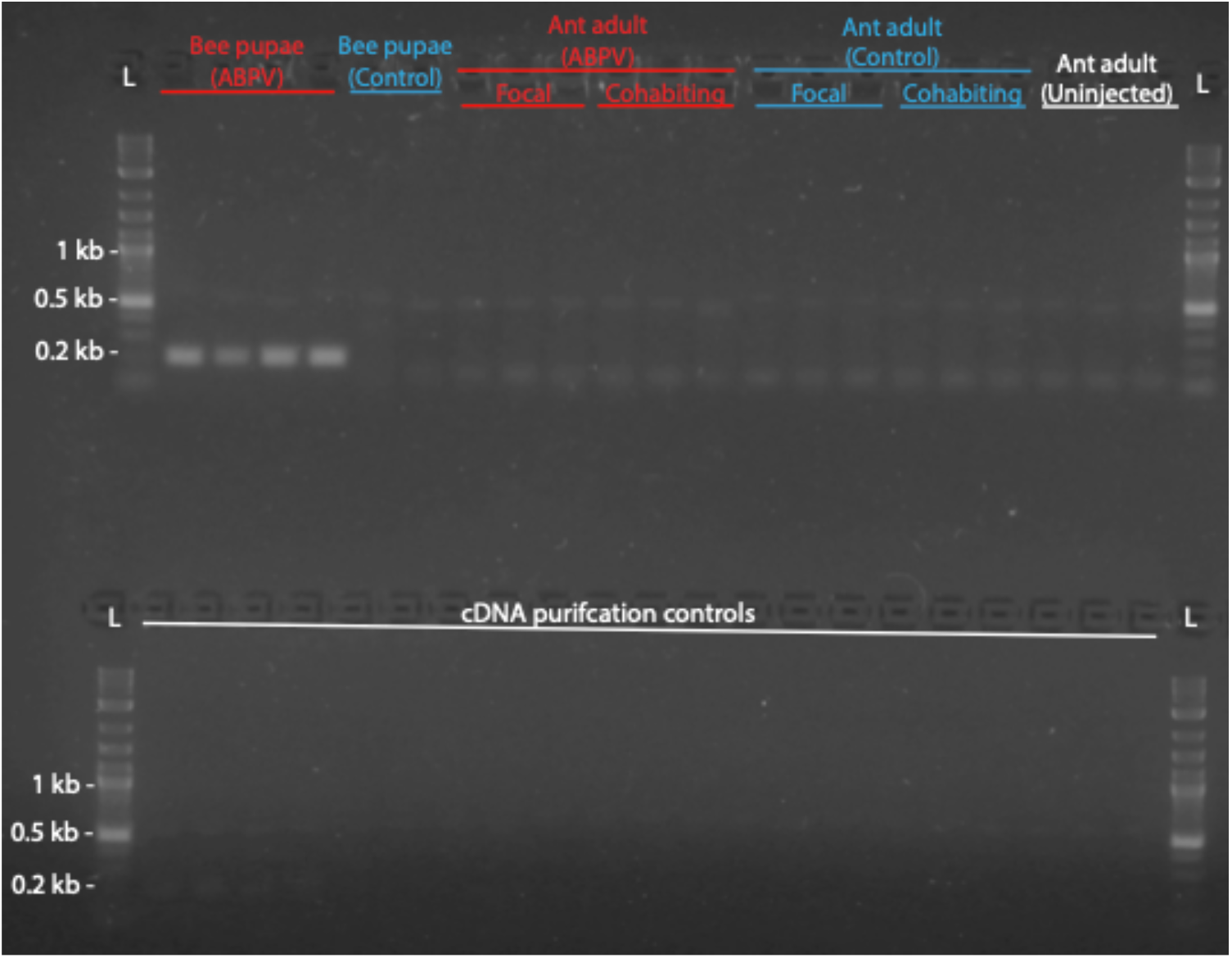
ABPV does not replicate in adult ants. Top row: ABPV-injected and control-injected bee pupae served as positive and negative controls, respectively. Red: ABPV-injected focals and cohabiting ants; blue: control-injected focals and cohabiting ants; white: uninjected. The expected PCR product size was 196 bp. L: DNA ladder. Bottom row: cDNA purification controls corresponding to the samples in the top row.

## 4. Discussion

Our study shows that injection of ABPV into *O. biroi* pupae and adults reduced host fitness by delaying development and decreasing survival. ABPV persisted over days in inoculated adults and pupae and efficiently spread to adults. Strikingly, uninoculated nestmates acquired ABPV titres comparable to those of directly inoculated ants. These results demonstrate that ABPV particles can persist and spread in ant colonies, and can impact host fitness, even in the absence of detectable viral replication.

The mortality observed in ABPV-injected *O. biroi* ads to the growing evidence that honey bee viruses such as ABPV can impair fitness in non-honey bee hosts, with outcomes depending on the inoculation methods and dosage. For example, black garden ants (*Lasius niger*) fed on ABPV-infected bee pupae for ten weeks showed impaired locomotion and reduced adult emergence (Schläppi et al., 2020). Similarly, IAPV, a close relative of ABPV (de Miranda et al., 2010), reduces host reproductive success and survival when fed to bumblebees at very high doses (2×10^7^ and 1×10^8^ particles) (Meeus et al., 2014; Piot et al., 2015; Wang et al., 2018), while injections of lower doses of IAPV (ca. 10^2^ particles) were sufficient to cause acute infection, paralysis, and mortality in bumblebee workers (Niu et al., 2014; Wang et al., 2016).

Despite strong host fitness impairment, we found no evidence of viral replication in *O. biroi* adults, as shown by stable or decreasing viral loads over time and a lack of detectable replicative viral strands. This suggests that fitness costs arise from host immunopathology (i.e., immune-mediated self-damage) rather than active infection (González-González & Wayne, 2020; Paludan et al., 2024), a common phenomenon across systems (Khan et al., 2017; Lazzaro & Tate, 2022; Rouse & Sehrawat, 2010; Scriba et al., 2024). For example, ABPV particles may trigger antiviral defences such as apoptosis and phagocytosis, but excessive apoptosis near vital organs can harm the host (Clarke & Clem, 2003; Nainu et al., 2017). Similarly, activation of the phenoloxidase cascade and the release of cytotoxic intermediates or reactive oxidative species confer broad antimicrobial activity, but can cause host tissue damage and early senescence (Dowling & Simmons, 2009; Mikonranta et al., 2014; Nappi & Christensen, 2005; Sadd & Siva-Jothy, 2006; Wu et al., 2012). In flies, overactivation or misregulation of the Imd pathway during early innate immune responses to invading pathogens or even benign gut microorganisms can lead to excessive immune responses that cause host mortality (Li et al., 2020; Paredes et al., 2011).

We speculate that the lack of ABPV replication in *O. biroi* is due to host specificity limiting viral infectivity and pathogenicity in a novel host (Doublet et al., 2025; Woolhouse et al., 2005). Host phylogeny strongly predicts infection outcomes, as shown for the *Drosophila* C virus and the Cricket paralysis virus, where changes in viral load correlate with relatedness between natural and novel hosts (Imrie et al., 2021; Longdon et al., 2011). Successful infection typically requires compatible host cell surface receptors for viral attachment and entry (Maginnis et al., 2018; Weaver et al., 2021) and ABPV may lack the receptor-binding domains required to interact with *O. biroi* cells. Additionally, viral host shifts are often constrained by host-specific physiological and immune barriers, requiring substantially higher inoculation doses to overcome immunity and initiate replication in a novel host (Gusachenko et al., 2020; Schläppi et al., 2020; Streicher et al., 2023; Tehel et al., 2022), a threshold potentially not reached in our experiments. Taken together with cross-species infection experiments in ants (Schläppi et al., 2020; Valizadeh et al., 2025) and bumblebees (Gusachenko et al., 2020; Tehel et al., 2022; Wang et al., 2018), our results indicate that although honey bee viruses can spill over into novel hosts and impair fitness, onwards interspecific transmission or spillback to honey bees is not guaranteed and greatly depends on exposure dosage and host-virus compatibility (Rivas Fontan et al., 2025; Tiritelli et al., 2024).

Adult ants cohabiting with ABPV-injected pupae or adults acquired detectable viral loads within one to three days. Despite an *ad libitum* diet, some ABPV-inoculated pupae disappeared or were found partially consumed, indicating brood cannibalism by workers, a common behaviour in *O. biroi*. In termites (Sun et al., 2013) and ants (Bizzell & Pull, 2024; Maák et al., 2020), brood cannibalism serves both hygienic (removal of infectious material) and nutritional (resource recycling) functions. Thus, consumption of infected nestmates or their carcasses by workers represents a plausible and efficient viral uptake route, consistent with findings in honey bees, where viruses can remain infectious in host tissues for 48 hours post-mortem (Payne et al., 2025) and where worker cannibalism of infected pupae is a major source of DWV spread within colonies (Posada-Florez et al., 2021). Cannibalism would also explain the unexpectedly high viral loads in adults cohabiting with pupae in our experiment, if entire infected pupae were consumed. However, we cannot rule out that viral spread may also have occurred by social contact, such as mutual grooming and brood care, or via environmental contamination of the shared nest space (Dalmon et al., 2021; Tehel et al., 2022).

Our work adds to the evidence that even without sustaining viral replication, ants can acquire and harbour bee viruses for extended periods (Schläppi et al., 2019; Valizadeh et al., 2025), potentially serving as incidental hosts or mechanical reservoirs that facilitate viral persistence in insect communities (Dobelmann et al., 2024). Positive-sense RNA viruses, such as ABPV, replicate rapidly and with high error rates, which can accelerate their adaptation to novel hosts (de Miranda et al., 2010). Consequently, even transient uptake by ants may promote low-level viral persistence that, under selective pressure, may give rise to variants capable of overcoming interspecific barriers and initiating host shifts, a process recognised as a key driver of emerging infectious diseases (Woolhouse et al., 2005). Environmental stressors, such as pesticide exposure (Al Naggar & Paxton, 2021; Schläppi et al., 2021) and climate change (Roberts et al., 2018), which affect host immune function and physiology, can potentially amplify the risk of adaptation to novel hosts and future spillover or spillback events, thereby shaping the epidemiology of bee pathogens in insect communities. Understanding these dynamics is critical for predicting viral emergence and designing strategies to safeguard pollinator health and enhance ecosystem resilience.

